# A Comparison of Peak Callers Used for DNase-seq Data

**DOI:** 10.1101/003608

**Authors:** Hashem Koohy, Thomas A. Down, Mikhail Spivakov, Tim Hubbard

## Abstract

Genome-wide profiling of open chromatin regions using DNase I and high-throughput sequencing (DNase-seq) is an increasingly popular approach for finding and studying regulatory elements. A variety of algorithms have been developed to identify regions of open chromatin from raw sequence-tag data, which has motivated us to assess and compare their performance.

In this study, four published, publicly available peak calling algorithms used for DNase-seq data analysis (F-seq, Hotspot, MACS and ZINBA) are assessed at a range of signal thresholds on two published DNase-seq datasets for three cell types. The results were benchmarked against an independent dataset of regulatory regions derived from ENCODE in vivo transcription factor binding data for each particular cell type. The level of overlap between peak regions reported by each algorithm and this ENCODE-derived reference set was used to assess sensitivity and specificity of the algorithms.

Our study suggests that F-seq has a slightly higher sensitivity than the next best algorithms. Hotspot and the ChIP-seq oriented method, MACS, both perform competitively when used with their default parameters. However the generic peak finder ZINBA appears to be less sensitive than the other three.

We also assess accuracy of each algorithm over a range of signal thresholds. In particular, we show that the accuracy of F-Seq can be considerably improved by using a threshold setting that is different from the default value.

## Introduction

Over the past decade, our ability to interrogate the features of the chromatin state has benefitted greatly from high-throughput sequencing (HTS) technologies. Genome-wide profiling of protein-DNA interactions has been made possible by Chromatin Immunoprecipitation coupled with high throughput sequencing (ChIP-seq) for a remarkable number of protein targets [1–3]. Similarly, HTS can be combined with the established DNase I hypersensitivity assay (DNase-seq) to profile open chromatin regions [4–7]. This approach has led to the detection of a total of nearly three million DNase I Hypersensitive Sites (DHS) across the human genome in about 140 different cell type [3, 8].

Probing the chromatin state using ChIP-seq and DNase-seq requires sophisticated data analysis pipelines once the sequence reads have been collected, but at their core, all analysis approaches involve gauging the significance of enrichment of short read tags in a given region relative to an expected background distribution. Algorithms used for this purpose are generally known as peak callers [6, 9].

Analysis of ChIP-seq data has received a great deal of attention and an enormous range of peak callers have been implemented [6, 10–14], benchmarked and extensively reviewed [2, 9–11, 15, 16]. However, DNase-seq has thus far received less attention and to the best of our knowledge there has been no systematic comparison of the performance of algorithms for calling DHSs from DNase-seq data. This places the end user in an uncertain situation, with little evidence to base decisions on as to which tools to use and with what parameter settings.

The properties of enriched regions vary greatly between different HTS-based chromatin interrogation technologies. For example, TF-ChIP experiments typically yield very sharp and punctate signals, while histone-ChIP for modifications such as H3K36me3 are much more broadly distributed. Signals from DNase-seq data, in turn, appear neither as sharp as those in TFBS ChIP-seq, nor as broad as in a typical histone modification ChIP [17, 18]. Therefore, peak callers that have been originally developed with ChIP-seq data in mind are usually not recommended for DNase-seq data, at least without additional parameter tuning [19].

To address this problem, a number of approaches have been presented. The Hotspot [7, 18] and F-Seq tools [20] have been implemented specifically for use with DNase-seq data (although F-Seq has also been used for ChIP-seq and FAIRE-seq data [21]). In contrast, Zero-Inflated Negative Bionomial Algorithm (ZINBA) [17] has been proposed as a generic tool for handling a variety of HTS data types including DNase-seq, FAIRE-seq, ChIP-seq and RNA-seq. Finally, several published studies have used the Model-based Analysis of ChIP-seq (MACS) peak caller [13] for the analysis of DNase-seq data [22]. As we will see, these tools are based on a diverse range of mathematical models, have different parameter spaces, and deal differently with the problem of background estimation.

In this paper, we compare the performance of the aforementioned four tools (all of which are open-sources and publicly available) on several DNase-seq datasets from the ENCODE project. The analysis has been performed on the chromosome 22 of the human genome GRCh37 assembly. The key aim of our analysis is to present a framework within which the user can decide which peak caller is more applicable to their data and whether or not the default signal threshold is appropriate in their case. In what follows, we first provide the reader with a brief overview of each of these peak callers and then present the results of our analyses.

## Results

### An Overview of Peak Callers

In this section we provide the reader with a brief description of each of the four tools used for bench-marking. More specifics about these algorithms including the version number, run time, the language in which they have been implemented and their original references are summarized in Table 1.

**Table 1.** DNase I Peak Callers Benchmarked in This Study.

| Algorithm | Version | CPU Time(sec.) | Max Memory(Processes) | Control Data | Mappability Data | Language | Refs |
| --- | --- | --- | --- | --- | --- | --- | --- |
| Hotspot | V3 | 6824 | 2288MB(10) | Not Required | Required | C++ | [7] |
| F-Seq | 1.84 | 1296 | 4304MB(3) | Not Required | Not Required | JAVA | [20] |
| MACS | macs2(2.0.10) | 562 | 1452MB(3) | Optional | Not Required | Python | [13] |
| ZINBA | zinba_2.01 | 412093 | 10318MB(10) | Optional | Required | R | [17] |

#### Hotspot

The Hotspot [7, 8] algorithm is the underlying algorithm used for the discovery of DHSs in the ENCODE project. The idea behind Hotspot is to gauge the enrichment of sequence tags in a region compared to the background distribution. Enrichment is measured as a *Z*–score, taking the binomial distribution of tag frequencies as the null model. Considering a small window of length 250bp centred in a larger window of length 50kb, the probability of each tag in the larger window hitting the small window is denoted as *p* which is defined as the ratio of the number of uniquely mappable tags in the smaller window to those in the larger window. (Note that *p* may differ in different regions because not all *k*–mers in a window can be aligned uniquely to the reference genome).

Assuming *n* tags hitting the smaller window and *N* tags hitting the larger window, the expected number can be calculated as *µ* = *N p*, the standard deviation as 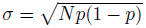, and the *Z*–score (that is then assigned to the small window) as 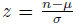. Using this method, each tag is assigned a *Z*–score which is equal to the *Z*–score of a small window centred at that tag position. Then a “hotspot” region is defined as a succession of tags having a *Z*–score above a specific threshold (assumed to equal two by default). Hotspot infers its final hotspots after two phases. Some highly enriched regions are detected as the first phase hotspots and the corresponding tags are filtered out from the set of short read tags. In the second phase, Hotspot tries to discover weaker but reproducible peaks that might have been overshadowed by the most enriched regions. Finally, the results of these two phases are combined and subjected to false discovery rate analysis. For this, Hotspot generates a set of random tags that is uniformly distributed over the mappable region of the genome. For a given *Z*–score threshold *T,* the *FDR* for the observed peaks centered at each tag with a threshold greater than or equal to *T* is defined as a ratio of the number of random tags with *Z*–scores greater than or equal to *T* to the number of observed tags falling within the same score range.

Hotspot is mainly programmed in *C*^++^, but the statistical analyses have been implemented in *R*. Some parts of the algorithm are also written in Python and as Unix shell scripts. The package depends on BEDOPS [23] and BEDTools [24].

A new implementation of Hotspot named “Dnase2hotspots” has been reported by Baek et al. [18]. The key difference between the two versions seems to be the merging of the two-pass detection in the original Hotspot algorithm into a single pass. At the time of our analyses, Dnase2hotspots required MATLAB for running, and was therefore excluded from the benchmarking. However, as this manuscript was at a late stage of revision, we learned that an updated version of Dnase2hotspots became available that no longer requires MATLAB (http://sourceforge.net/projects/dnase2hotspots/).

#### F-Seq

F-Seq [20] was developed with the aim of summarising DNase-seq data over genomic regions. The authors identified problems with histogram based-peak calling algorithms, in which the enrichment of tags is measured across equal-sized bins. Such algorithms suffer from boundary effects and difficulties in selecting bin widths.

In F-Seq, it is assumed that *n* short tags {*x*_*i*_} are independently and identically distributed along the chromosome i.e. 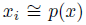 such that the probability density function is inferred as: 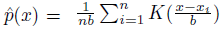 in which *b* is the bandwidth parameter to control the smoothness and *K*() is a Gaussian kernel function. Although this algorithm was initially developed for DNase-seq data, it has also been used for ChIP-seq peak detection [20].

#### ZINBA

ZINBA [17] is a generic algorithm for genome-wide detection of enrichment in short-read data that was proposed for the analysis of a broad range of genomic enrichment datasets. ZINBA first divides each chromosome into small non-overlapping windows (250bp by default) based on the number of reads. These read count values, alongside other covariates including G/C content, mappablility scores, copy number variation and an estimation of background distribution make up the parameters of a mixture regression model. This model then assigns each region into one of three classes: enriched, background, or zero (windows for which no read is assigned due to insufficient sequencing coverage). The relationship between the covariates and the signal for various experimental data is then inferred through an Expectation Maximisation-based implementation of a mixture regression model. ZINBA is supplied as an *R* package.

#### MACS

MACS [13] is one of the most popular peak callers for ChIP-seq data [14] that has recently been used for DNase-seq [22]. As a ChIP-seq tool, MACS has been reviewed and benchmarked in a number of studies [9,10,12]. The key advantage of MACS compared to previous peak callers is that it models the shift size of tags and can also allow for local biases in sequencability and mappability through a dynamic Poisson background model. MACS is written in Python and can be run with or without an input control dataset. The only required input for this model is a set of short read tag alignments.

### The Sensitivity and Specificity of the Peak Callers

To systematically evaluate the performance of Hotspot, F-Seq, MACS and ZINBA, we ran them on the publicly available DNase-seq data sets for K562, GM12878 and HelaS3 cell type over human chromosome 22 [8] (see Methods for the availability of these data sets). A visual inspection of peaks generated by these peak callers (Figure 1A) at their default signal threshold showed that their were not fully consistent. In particular, it can be seen that while some regions of strong enrichment were consistently detected, there was a significant variation in the detection of weaker regions, as well as in the sizes of the recovered DHS peaks. To our surprise, only *∼* 11.5% of the reference set (at the base pair level) were consistently detected by all four tools (8% in K562, 13% in GM12878 and 14% in HeLaS3, respectively). Overall, peaks detected by at least one tool spanned on average 41% of the reference set (30% in K562, 48% in GM12878 and 46% in HeLaS3, respectively). This is likely due to a combination of factors, including the genuine mapping of some TF binding sites in the reference set outside of regions of increased chromatin accessibility, some “true” DHSs missed by the DNase-seq protocol and the false-negative rates of the peak detection tools themselves. The base-pair overlap of the peak regions detected by each algorithm in the three cell lines is shown in Figure 1B. Significant differences were also observed in the running times of the algorithms, with ZINBA taking on average 370x longer and using 4.5x more memory than the rest (Table 1).

**Figure 1.**
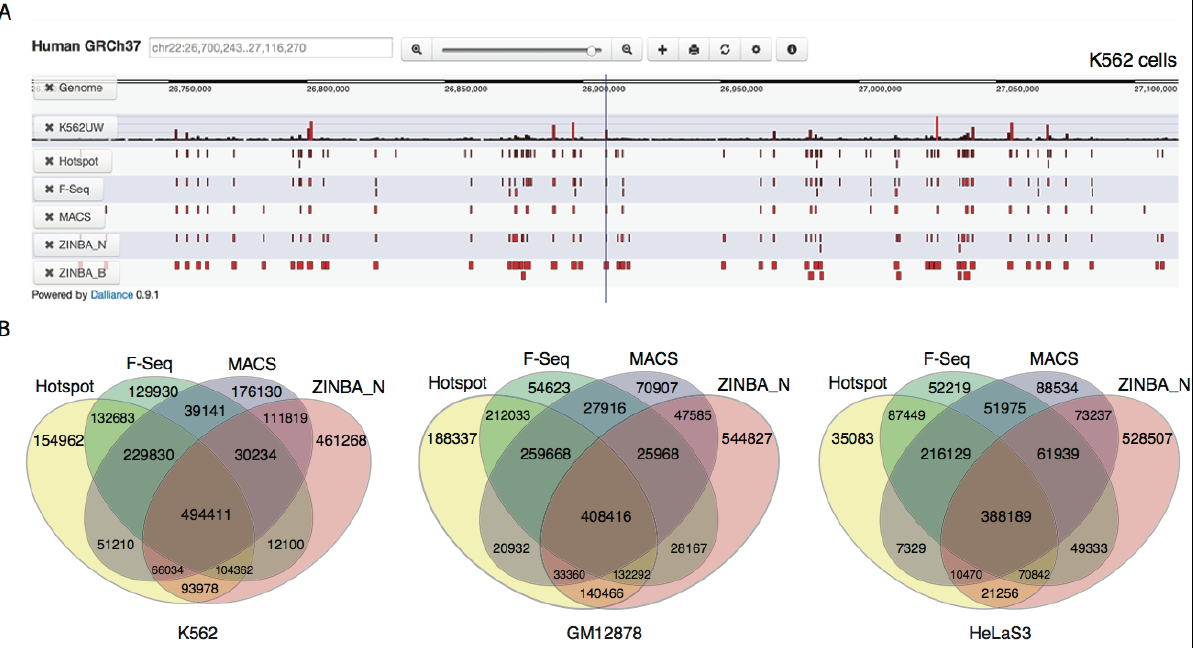
Comparison of the Four Peak Callers in a Representitive Genomic Region. (A) A screenshot from Dalliance [29] showing peaks called by the four peak callers in about 400kb of chromosome 22 in K562 cells. The first row in this figure labelled as ‘K562UW’ illustrates the distribution of short read tags of K562 (replicate 1) from University of Washington (see Methods for full details). The following rows show the statistically significant regions (peaks) according to each of the algorithms with their default signal thresholds. (B) Overlap between peaks called by each algorithm. Venn diagrams showing the overlap between peaks called by each of the four algorithms using their default parameters in K562 cells (left), GM12878 cells (middle) and HeLaS3 cells (right). The numbers correspond to the number of basepairs called.

We then ran each of these four tools over a range of signal thresholds and compared the peaks detected by each algorithm at each threshold level to the “reference sets” of regulatory regions. These sets were generated by pooling the ChIP binding profiles of multiple transcription factors (TFs) in each of the three cell types (the ChIP data was produced by ENCODE [15], see also Supplementary Data (Files GM12878, K562 and HeLaS3) for the list of TFs used). Using TF-binding profiles to produce the reference set has been motivated by the fact that the majority of TF binding sites map to regions of increased chromatin accessibility that are detectable as DNase hypersensitive sites [5, 8]. Although our reference set is inevitably incomplete, since the ChIP data is only available for a subset of TFs, it still allows us to robustly assess the relative performance of the DHS-calling algorithms (as used previously in [25]).

For each of the four algorithms, we estimated the sensitivity (expressed in the terms of the True Positive Rate, *TPR*) and specificity (expressed as 1 – *FDR*, False Discovery Rate) from the degree of the overlap (at base pair level) of their respective DNase I peaks at each signal threshold with each of the reference sets. This approach is presented in more detail in the Methods section. The sensitivity-specificity analysis revealed further substantial differences between the peak finders (Figure 2). In particular, we found ZINBA to underperform all other tested tools in terms of both *TPR* and *FDR*. Its “narrow peaks” output (ZINBA_N) showed the lowest *FDR* among all algorithms, but also the lowest *TPR*, meaning that ZINBA_N may miss many true DHSs. On the other hand, ZINBA’s “broad peaks” output (ZINBA_B) still had a relatively low *TPR* but also showed the highest *FDR*, meaning that its broad peaks showed a poorer overlap with the reference set compared to the other three peak callers.

**Figure 2.**
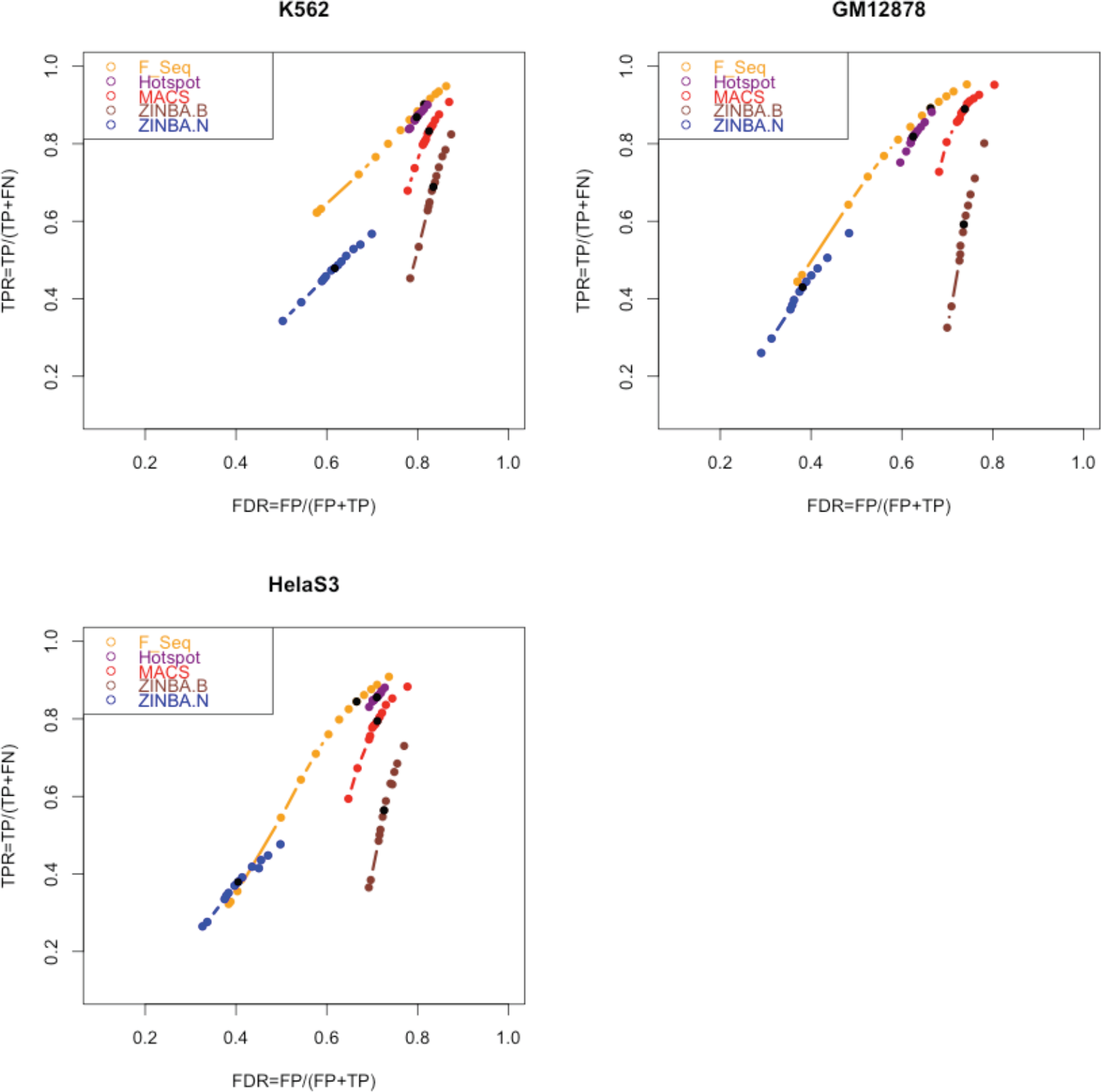
Comparison of the Peak Calling Algorithms Based on Estimated True Positive and False Discovery Rates. Each algorithm was run over 13 values of a parameter that controls the false discovery. These values for Hotspot, MACS and ZINBA range from 0.001 to 0.2 and for F-Seq it ranges from 0.001 up to 6 (see methods for more details). For each value the overlap between the calls and the “reference set of regulatory regions” for that cell type was measured. The black dots show the default value for each algorithm.

Both the *TPR* and *FDR* of the other three peak callers (Hotspot, F-Seq and MACS) segregated by nearly 10% on the data from the GM12878 cell type. As we can see from Figure 2, in this cell type, F-Seq showed the highest *TPR* and Hotspot showed the best (lowest) *FDR*. More similar *FDR* and *TPR* values were observed in the other two cell types, with both F-Seq and Hotspot having only slightly lower *TPR* and higher *FDR* compared to MACS (Figure 2).

We asked if the relative performance of the algorithms is affected by the choice of a specific DNase-seq protocol. Currently, there are two DNase-seq protocols commonly used by the community: the “end capture” protocol [26] and the “double hit” protocol [27]. While this study so far focused on the “double hit” protocol, we also evaluated the performance of the algorithms with the “end capture” protocol using the ENCODE data for the K562 cell type [8]. However, we found the relative performance of the algorithms to remain generally consistent across the two protocols (Figure S1).

Overall these results suggest that F-Seq, Hotspot and MACS generally outperform ZINBA with DNase-seq data in terms of both specificity and sensitivity, with the F-Seq algorithm showing the best performance of all four algorithms tested.

### Comparison of the Summary Statistics of the Detected Peaks

We next sought to evaluate how the differences in the performance of the four algorithms are reflected in the summary statistics of the respective peaks. As shown in Figures 3 and 4, peaks detected by the four algorithms vary both in the total number and their length distributions. In particular, MACS produced the smallest number of peaks compared to the other three algorithms, followed by ZINBA (for which the numbers of broad and narrow peaks were equal). The peaks from F-Seq and Hotspot outnumbered both MACS and ZINBA peaks, with either F-Seq or Hotspot yielding the highest number depending on the cell type.

**Figure 3.**
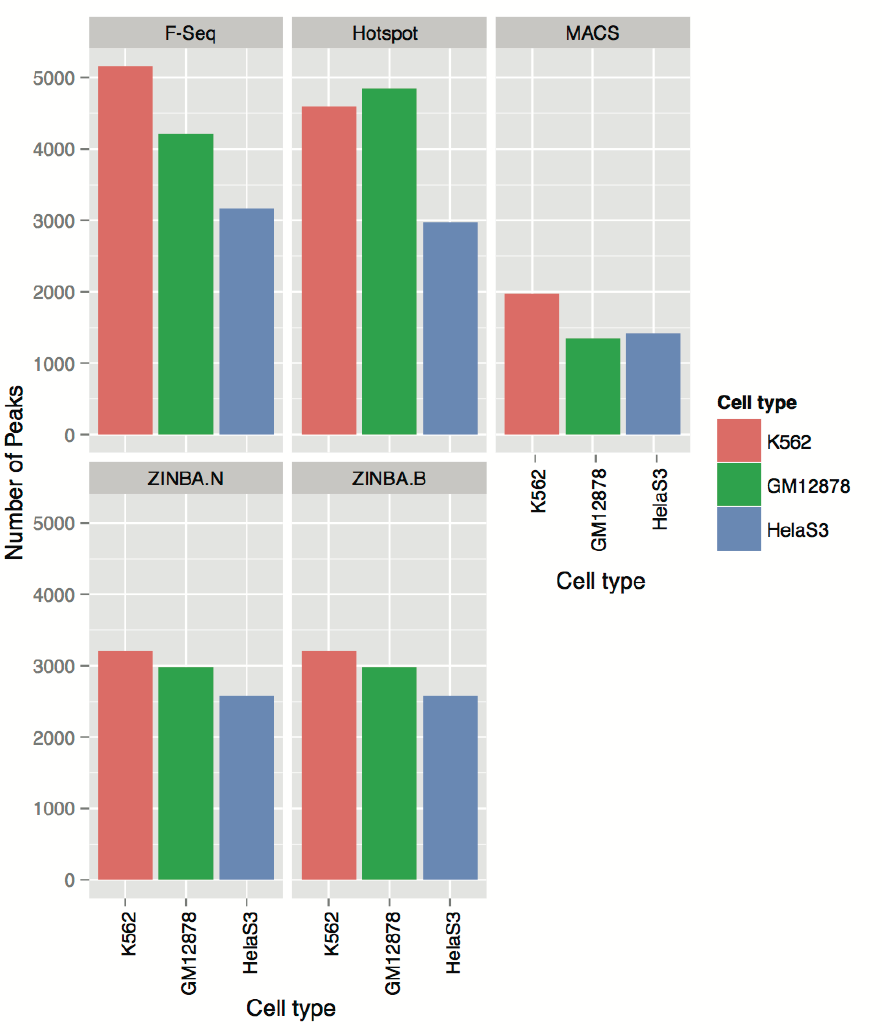
Number of Peaks Detected by Each Peak Caller Using Their Default Parameters. The number of peaks obtained by each algorithm at their default signal threshold.

**Figure 4.**
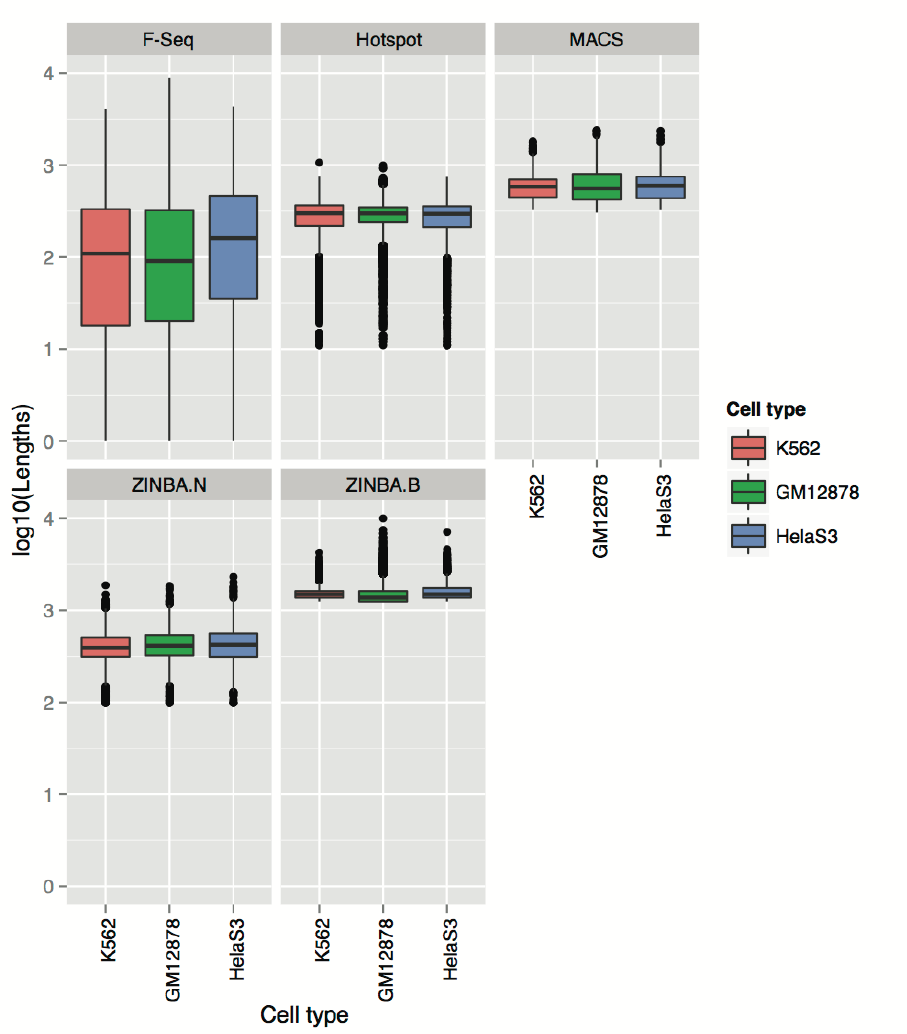
Distribution of Lengths Depending on Peak Callers and Their Parameter Settings. Distribution of peak lengths found by each of the algorithms, when ran with their default parameters, are compared between cell types. ZINBA.N and ZINBA.B represent narrow peaks and broad peaks (respectively) obtained from ZINBA.

ZINBA’s broad peaks were on average the longest compared to all other datasets, ranging from 1kb to 10kb (Figure 4). These were followed, sequentially, by MACS peaks (with a median of around 2700bp over all three cell type), ZINBA narrow peaks and Hotspot peaks (median length 2.5kb). F-Seq peaks were on average the shortest, with a median of 2kb but notably, they showed a considerably higher variance of peak lengths (Figure 4).

These differences prompted us to look at the overall peak coverage produced by each algorithm, which we defined as the ratio of the number of base pairs covered by the peaks to the length of the chromosome. Note that chromosome 22 has an active arm of about 35Mb. It can be seen from Figure 5, with the exception of ZINBA.B (broad) peaks showing an appreciably higher coverage than the rest, the peaks from all four algorithms (including ZINBA’s narrow peaks) showed a comparable coverage. On average, MACS showed the lowest coverage and ZINBA.N showed the greatest coverage among the narrow peaks of algorithms. The highest spread of coverage (1.35%) was observed in GM12878 cells, between ZINBA.N (3.88%) and MACS (2.53%). The lowest spread of 0.6% was observed in K562 cells. The similarity in the peak coverage produced by the four algorithms at their respective default parameter settings suggests that these settings were generally appropriate for a relative evaluation of the tools’ performance.

**Figure 5.**
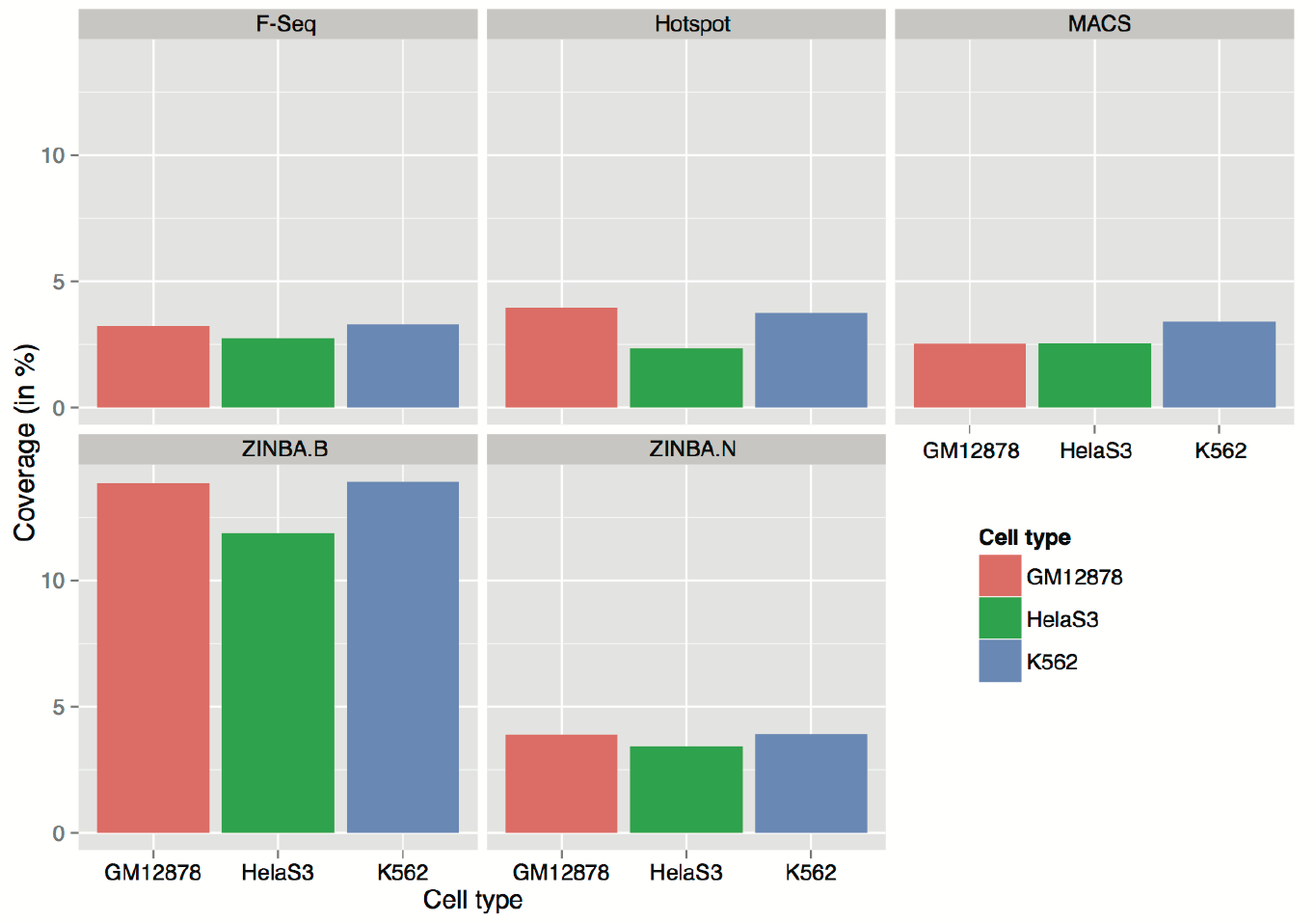
Coverage of Peaks Detected by Each Peak Caller Using Their Default Parameters. Illustrated here is the percentage of chromosome 22 covered by peaks from each peak caller over three cell type.

### Effects of Algorithm-specific Parameters

So far, we have compared the algorithms’ performance across the range of a single parameter that was common to all four peak callers: the overall signal threshold for making a peak call. Although a number of additional, mostly algorithm-specific, parameters exist, we kept them at their default values. A comprehensive evaluation of the peak callers over their full parameter spaces is challenging due to the algorithm-specificity of some parameters and also to the extensive number of parameter combinations. Some of these parameters, however, are unlikely to affect the sensitivity or specificity of the algorithms, as they are concerned either with other data types (eg ChIP-seq) and/or file formats. For example, in MACS one may see “–broad” and “–call-summits” for data type, “-g” for genome size and “-f” for file format. However, a number of tunable parameters, in particular in Hotspot and ZINBA seemed to affect the key parts of the respective algorithms, prompting us to ask whether they have a significant effect on the results.

For Hotspot, we evaluated the effects of the *z*–score and the merging size threshold. As shown in Figure S3, the distribution of peaks’ lengths is nearly indistinguishable when merging peaks closer than 150bp (default) or not merging them at all. Similarly, we found that the performance of the Hotspot remains almost invariable at a range of *z*–scores (*z* = 1, 2, 3, 4; Figure S4).

For ZINBA, we assessed the effect of the number of hits per read allowed during mapping process (“athreshold”), and of average fragment library length (“extension”) on its performance. As can be seen from Figure S5, peak coverage remained insensitive to varying the “athreshold” parameter. In contrast, increasing the “extension” parameter from the default resulted in the peak coverage increasing beyond the range observed for all other peak callers.

In conclusion, we found no evidence that adjusting the algorithm-specific parameters of Hotspot and ZINBA leads to improved performance compared to their default parameter settings.

### Adjusting the Default Signal Threshold Setting Improves the Performance of F-Seq

As a final step in our analyses, we set out to determine the peak signal threshold settings that ensure an optimal tradeoff between sensitivity and specificity. To this end, we expressed the sensitivity and specificity data for each peak caller generated over a range of signal thresholds (described above and shown in Figure 2) in terms of the *F*–score metric which is commonly used in information retrieval. The *F*–score combines both the sensitivity and specificity such that the higher *F*–score values indicate a more optimal performance (see Methods and also [25]). The relative contribution of sensitivity and specificity is weighted by the *β* parameter that we assumed to be 0.5 to place a higher emphasis on specificity over sensitivity (see Methods for more detail).

In Figure 6, we plotted the *F*–scores corresponding to a range of peak thresholds for each of the tools. As can be seen, F-Seq showed an improved performance when its signal threshold (defined by the “standard deviation threshold” parameter) was reduced from the default value of 4 to a value between 2 and 3. In contrast, Hotspot performance remained largely unchanged over the range of its threshold parameter. For MACS, the default threshold settings seemed optimal. ZINBA on the other hand, showed continuously decreasing *F*–scores with increasing threshold, suggesting no clear-cut optimal threshold setting.

**Figure 6.**
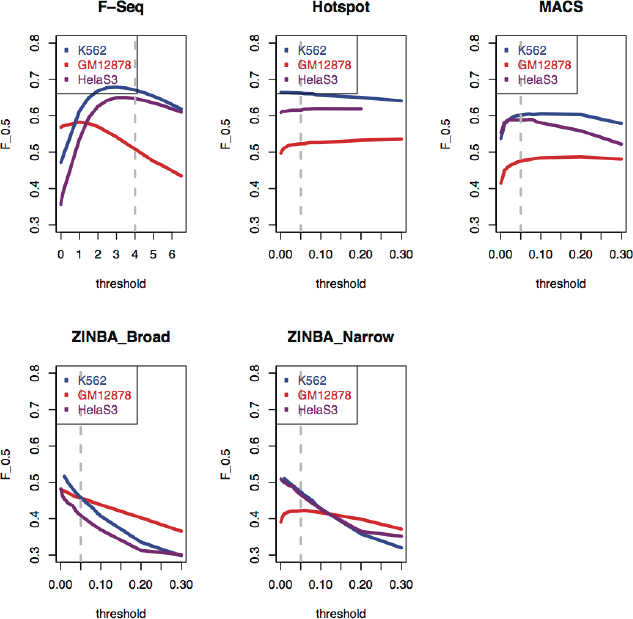
F Scores of Algorithms Over Three Cell Types From the “Double Hit” Protocol. Each algorithm was evaluated to gauge the enrichment of short read tags in each of the three cell types obtained from University of Washington “double hit” protocol [27]. The overlap of peaks from each of the cell types was measured against the cell type’s “reference set of regulatory regions”. The accuracy of each algorithm was defined as the value of the F score (see Methods for more details) by running it over a range of thresholds. The dashed vertical grey line depicts the value of F score when the algorithm is run with its default parameter. Note that Hotspot failed when ran with *FDR* = 0.3 for HelaS3 cell type and therefore its corresponding curve is shorter by one data point.

In conclusion, while Hotspot and MACS showed a near-optimal performance at the default signal threshold settings, the performance of F-Seq can be further improved by reducing the threshold parameter.

## Discussion

In this study, four open-source peak callers proposed for the analysis of DNase-seq data were bench-marked and briefly reviewed. Our results showed that there is, in fact, a considerable discrepancy in the tools’ performance. Of the four peak callers, F-Seq showed the best performance with DNase-seq data, particularly when run with a signal threshold level slightly lower than default. Both Hotspot and MACS also showed appreciable performance, only slightly lagging behind F-Seq in both sensitivity and specificity. In contrast, and despite its reported performance with RNA-seq, ChIP-seq and FAIRE-seq data [17], ZINBA showed to be less suitable for DNase-seq data analysis, both in terms of specificity, sensitivity and the computational time. To the best of our knowledge, this peak caller has not been used with DNase-seq in any published studies.

Although both ChIP-seq and DNase-seq experiments generate short-read tags, there exist a number of differences between these data types that caution against the application of ChIP-seq peak callers to DNase-seq data, at least without re-tuning their parameters. The key differences include: a) ChIP-seq data usually shows a higher signal-to-noise ratio compared to DNase-seq, making ChIP-seq peaks easier to detect; b) ChIP-seq data, unlike DNase-seq data, are strand-specific with a shift in the signal between strands; c) as the general hallmarks of open chromatin regions, DHSs may cover wider regions, spanning the binding positions of different regulators and differentially modified histones; therefore DHSs vary more broadly in length compared to typical ChIP-seq peaks [9, 19]. Taking these differences into account, one may conclude that the ChIP-seq-oriented peak caller MACS performs relatively well for DNase-seq data.

In our analyses we benchmarked the performance of each algorithm against a “reference set” of regulatory regions, generated from the union of multiple TF-binding profiles from ENCODE. This allowed us to compare the results of the peak callers with a “standard” that is based on a different type of experimental data and that is analysed using a different set of tools. It must be noted that, despite the large number of TFs used, our “reference set” is necessarily incomplete and may have its own inherent biases. It seems unlikely that these biases would selectively favour the performance of some DNase-seq algorithms over others. The continued expansion of the range of TFs profiled by ChIP will make it possible to further improve the precision of such reference sets in the future.

Furthermore, we recently showed that DNase I has DNA binding preferences [25] that potentially present a source of bias in DHS detection. This largely unexpected property of the DNase I enzyme is currently unaccounted for by any peak caller. There may therefore be scope for a new generation of DHS peak calling algorithms taking this factor into account.

Primarily due to ZINBA’s extended run time (see Table 1), benchmarking was limited to chromosome 22. To the best our knowledge, chromosome 22 is a representative part of the human genome, at least with respect to the density and distribution of TF ChIP peaks and DHSs. It is therefore expected that the benchmarking results obtained on chromosome 22 are applicable genome-wide.

Finally, it is worth mentioning that in addition to the quality of peak calling per se, factors such as documentation and the overall user friendliness may play a role in the choice of DNase-seq analysis software, particularly by experimental biologists. To this end, F-Seq, MACS and ZINBA are published and well-documented (see [20], [13] and [17]). Hotspot has been partly described in [7], but its source code and some more documentation are available at http://www.uwencode.org/proj/hotspot-ptih/.

DNase-seq is gaining popularity as a genome-wide chromatin accessibility analysis method, and its applications have led to new insights into genome function and variation [8, 28]. Robust peak detection on these data is therefore instrumental to the research community, particularly when it is provided by publicly available, well-documented and user-friendly software that can be easily used in any lab.

## Materials and Methods

The performance of four peak calling algorithms was compared over a range of the false discovery rate thresholds for Hotspot, MACS and ZINBA and a range of the standard deviation threshold for F-Seq. Each of the methods was used on the DNase-seq short-read data from three cell type (K562, GM12878 and HelaS3) that was obtained from the ENCODE project [8, 26]. We assessed the performance of these methods by comparing the peaks reported from each of these algorithms to the “reference sets of regulatory regions” generated from a union of peaks from a set of transcription-factor binding ChIP experiments for each of the three cell type. Our analyses were restricted to chromosome 22, primarily due to the very significant compute times taken by ZINBA. All data in this study was mapped to the GRCh37 (hg19) human genome assembly. All computations were run on an Intel(R) Xeon(R) CPU *E*5440 @ 2.83*GHz*, with 6GiB of RAM.

Our experimental design was as follows:

### Step 1: Input files

We downloaded University of Washington DNase I short read tags for K562, GM12878 and HelaS3 from http://hgdownload.cse.ucsc.edu/goldenPath/hg19/encodeDCC/wgEncodeUwDnase/ and for Duke University from http://hgdownload.cse.ucsc.edu/goldenPath/hg19/encodeDCC/wgEncodeOpenChromDnase/ as BAM files which are labeled as wgEncodeUwDnaseK562AlnRep1.bam, wgEncodeUwDnaseGm12878AlnRep1.bam and wgEncodeUwDnaseHelas3AlnRep1.bam. The number of short read tags mapped to chromosome 22 were 434301, 426770 and 255489 respectively for K562, GM12878 and HelaS3.

### Step 2: Running peak callers at different thresholds

We ran Hotspot, F-Seq, ZINBA and MACS with the aligned datasets listed above (either directly from the BAM files or converted to BED format if required) with the following thresholds:

**Hotspot**: Keeping all other parameters in Hotspot as their defaults, we tried the *FDR* threshold with values equal to *0.001, 0.005, 0.01, 0.02, 0.03, 0.04, 0.05, 0.06, 0.07, 0.08, 0.09, 0.1, 0.2, 0.3*.
**F-Seq**: Although there isn’t a parameter defined in F-Seq to directly control *FDR*, the standard deviation threshold *t* defined in F-Seq has an inverse correlation with *FDR* [20]. The default *t* in F-Seq is equal to 4. In this analysis we therefore ran it with an *t* equal to *0.001, 0.005, 0.05, 0.1, 0.5, 1, 1.5, 2, 2.5, 3, 3.5, 4, 4.5, 6* The feature length parameter (representing the bandwidth) was equal to 600bp by default.
**MACS**: The parameter controlling the *FDR* in MACS is called q-value and its default is 0.05. In our analysis we ran it with a *q* equal to *0.001, 0.005, 0.01, 0.02, 0.03, 0.04, 0.05, 0.06, 0.07, 0.08, 0.09, 0.1, 0.2, 0.3*.
**ZINBA**: In ZINBA the signal threshold controlling the *FDR* is called “threshold”, with a default value of 0.05. In this study we ran it with thresholds of *0.001, 0.005, 0.01, 0.02, 0.03, 0.04, 0.05, 0.06, 0.07, 0.08, 0.09, 0.1, 0.1, 0.2, 0.3*. Inspired by the developers’ demonstration for the FAIRE-seq data, we set *numProc* = *5* and *extension* = *150*.

### Step 3: Making a reference set of regulatory regions

For each of the cell types K562, GM12878 and HelaS3, we downloaded the narrow peaks of 99, 53 and 56 TFBSs respectively from the ENCODE project repository at http://hgdownload.cse.ucsc.edu/goldenPath/hg19/encodeDCC/wgEncodeSydhTfbs/ (these were all the available TFBSs as SydhTfbs for these three cell type) See Files S1, S2 and S3). Then we computed the union of TFBSs (using [23]) at each cell type and took it as our reference set of regulatory regions specific for that cell type.

### Step 4: Measuring the performance of the algorithms

We defined the overlap (at base pair level) between peak calls of each algorithm at each threshold and our reference set of regulatory regions as a metric for measuring the performance of each of the algorithms. More precisely, for each algorithm and for each threshold, the True Positive Rate (also known as sensitivity) was defined as 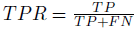 which is in fact the ratio of the number of correctly predicted base pairs to the number of base pairs in the union of TF set. Similarly, the False Discovery Rate was defined as 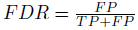, which is the ratio of the number of falsely found bases as peaks to the whole set of peaks found. The reader should take care to distinguish between the *FDR* that we have defined here and the false discovery threshold parameter defined in each of Hotspot, MACS and ZINBA algorithms.

The specificity (or precision) in this context was defined as 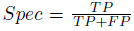 and the sensitivity was defined as 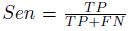, which is sometimes called “recall”. For each experiment the *TPR* was plotted against *FDR*.

Common to information retrieval, the overall performance of algorithms was defined as an *F*–measure:

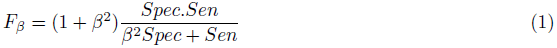

As can be seen from this equation, *F_β_* assigns *β* times as much weight (or importance) to sensitivity as specificity. Normally, in situations where both specificity and sensitivity are of equal importance, *β* is set to 1, and the score is known as *F*_1_ or as the “harmonic mean”. In our analysis, however, because of incompleteness of our reference data set (TFs), we used *F*_0.5_ to put more emphasis on specificity than sensitivity. Our choice of *β* reflects our prior belief about the incompleteness of the reference set. Using other reasonable values of *β* does not significantly affect our conclusions about the relative performance of the algorithms. For example, Figure S6 shows the results from Figure 6, but assuming *β* = 1 (*i.e.* an equal emphasis on specificity and sensitivity), instead of *β* = 0.5.

## Acknowledgments

We thank Alistair Rust, Ignacio Vazques Garcia and Daniel Bolland for their helpful comments. Advice from Naim Rashid and Paul Giresi on generating mappability data is appreciated.

---

**Figure S1.**
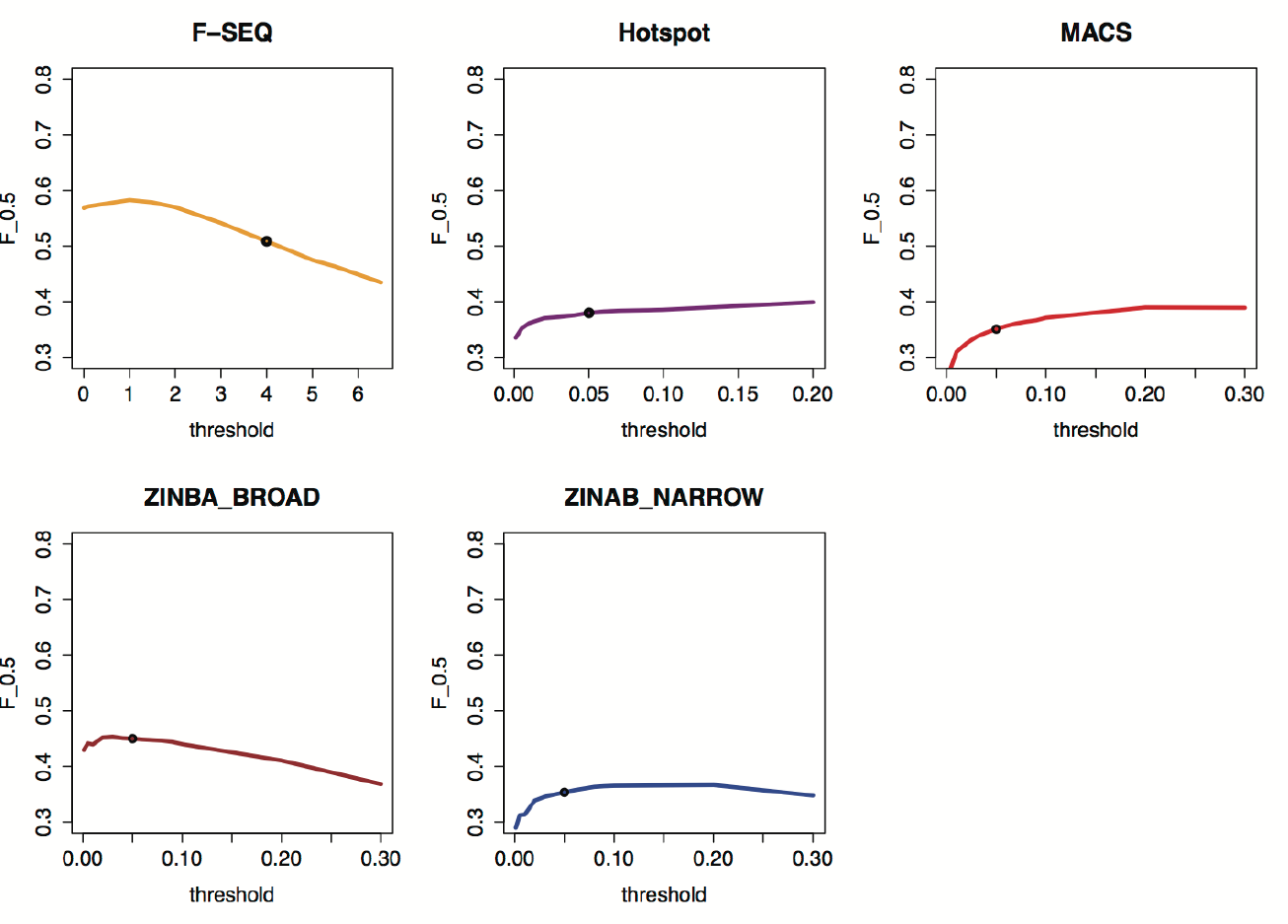
Performance of Algorithms Over One Cell Type From the “End Capture” Protocol. Similar to Figure 6, the performance of each algorithm was evaluated using GM12878 cell type obtained from Duke University “end capture” protocol [26].

**Figure S2.**
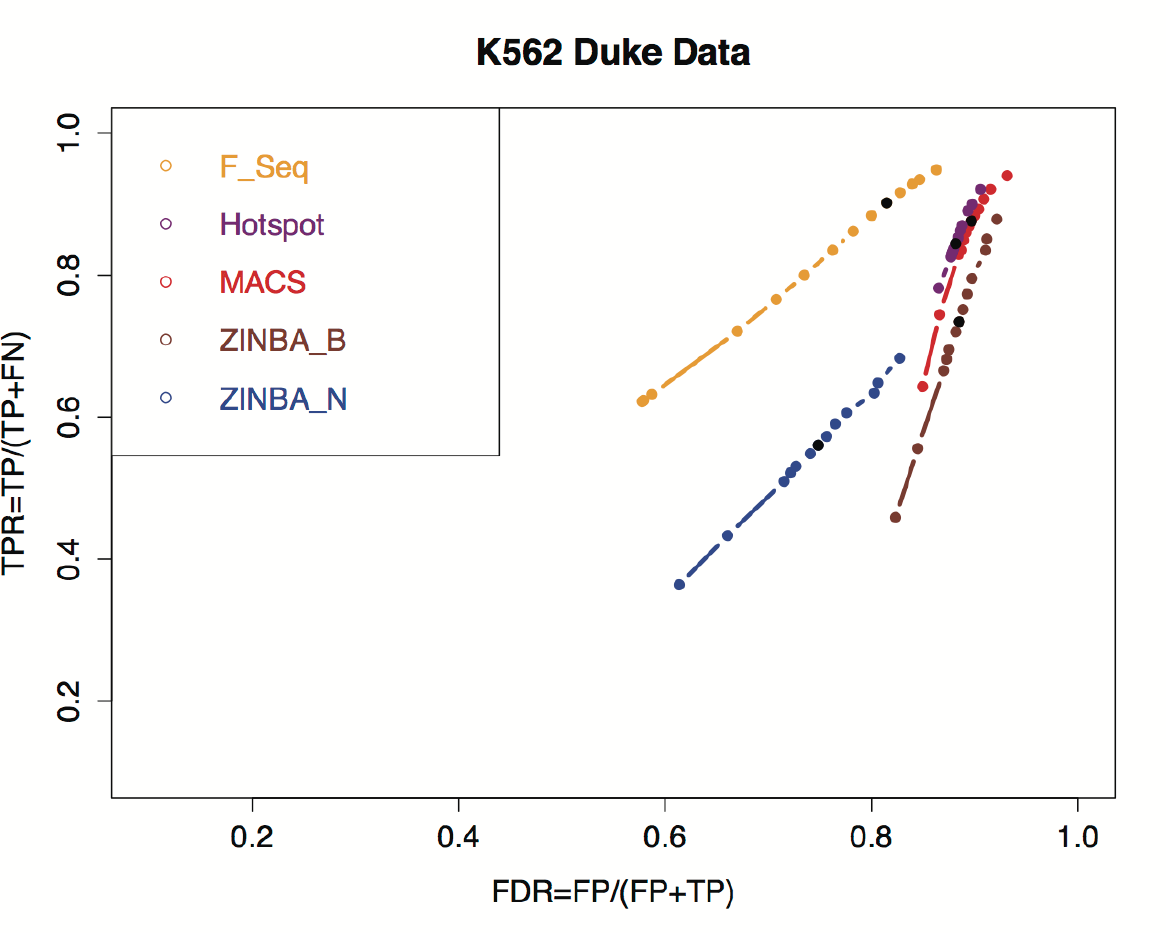
Comparison of TPR and FDR of Peak Callers with “End Capture” Data. Depicted here is the result of our *TPR – FDR* comparison of four algorithms over data obtained from Duke University end capture protocol.

**Figure S3.**
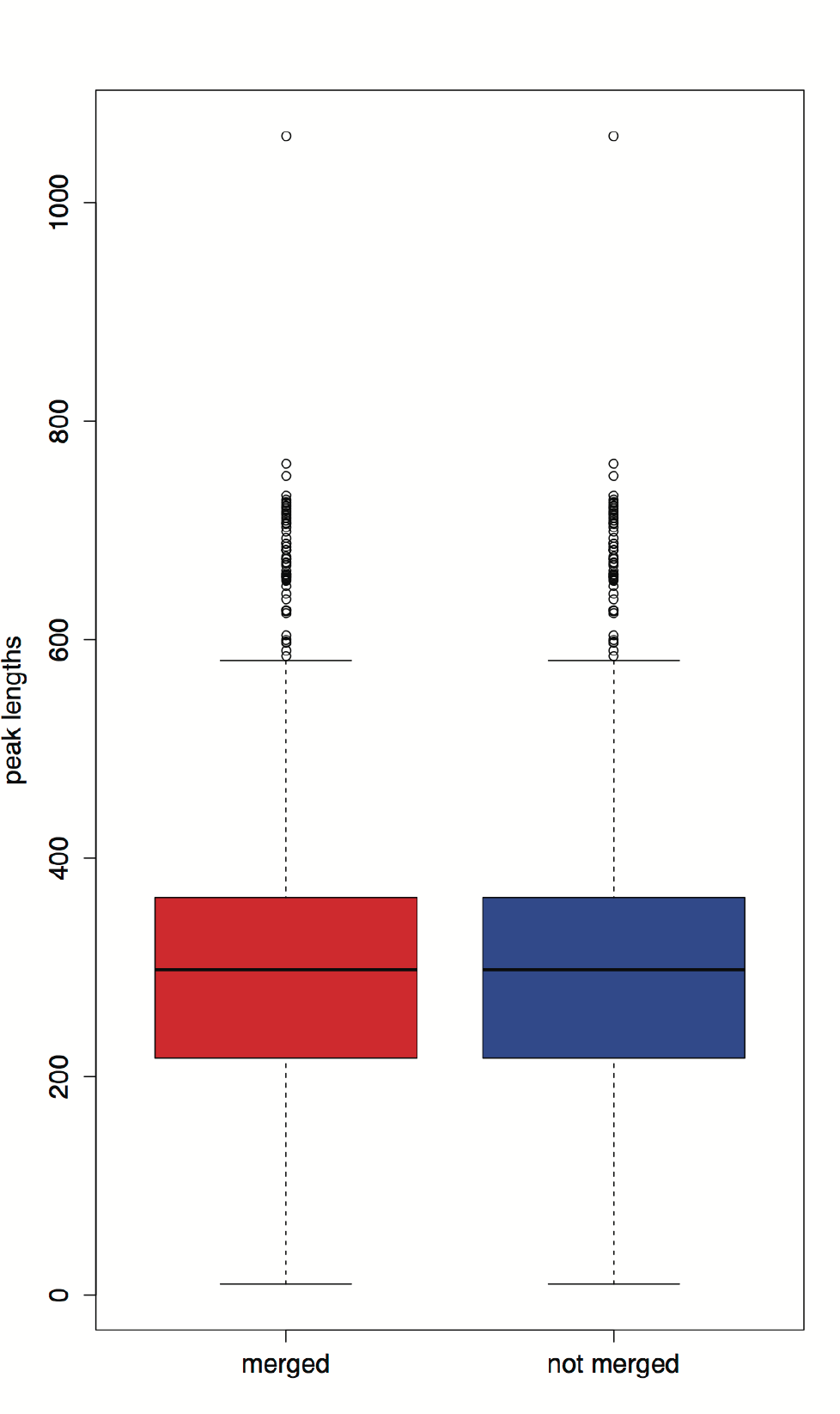
Effect of Hotspot “merge” Parameter on the Distribution of Peak Lengths. Distribution of Hotspot peak length merged (default: peaks closer than 150bp are merged) versus not merged in UW K562 cells.

**Figure S4.**
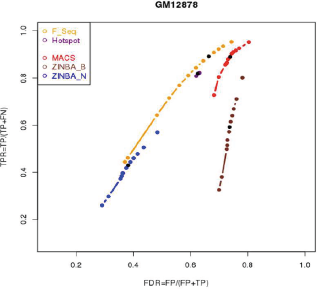
Effect of Hotspot “zscore” Parameter on its Performance. Hotspot was run at a range of z-score threshold ranging from 0.5 to 4 and all other parameters were kept as default. The other three algorithms were also run at a range of signal threshold (as described in main text).

**Figure S5.**
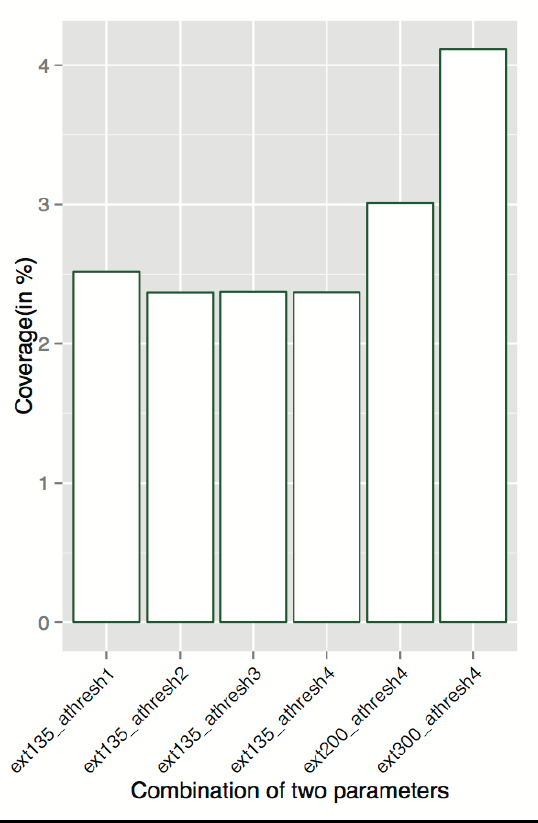
Effect of the Number of Hits and Extension on ZINBA Coverage. Depicted here is the coverage (as defined in main text) of ZINBA when run at various combinations of number of hits per read known as “athreshold”(run at values equal to 1, 2, 3, 4) and the average of fragment lengths known as “extension”(run at values equal to 135, 200 and 300bp).

**Figure S6.**
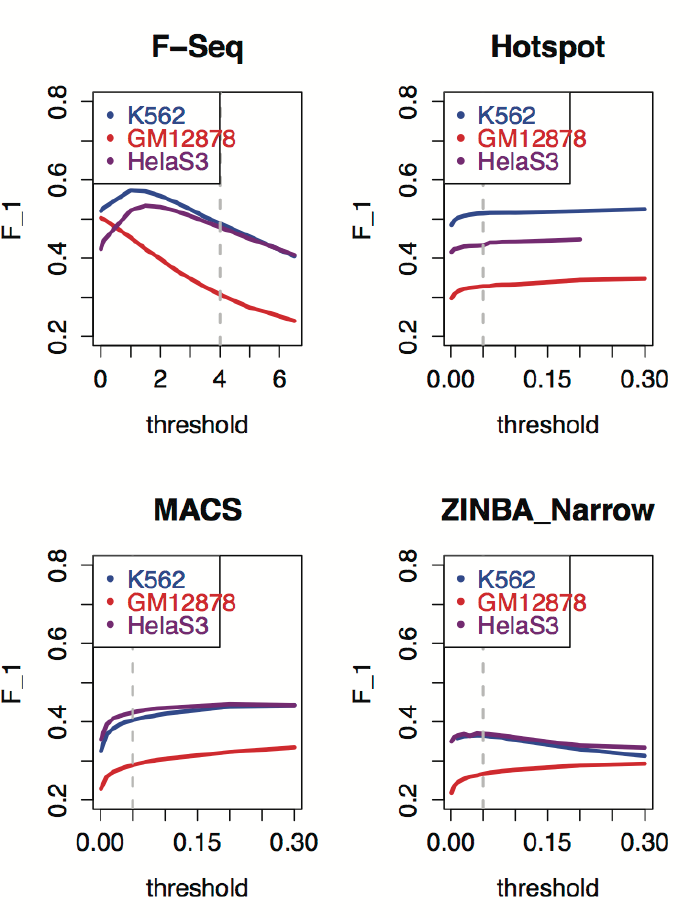
The F-scores of the Algorithms Across the Three Cell Types Assuming *β* = 1. Illustrated here is the data shown in Figure 6, but computed assuming the *β* parameter equal to 1, which corresponds to same weight associated with both sensitivity and specificity. The vertical dash lines show the default threshold values in each algorithm.

